# scDTL: single-cell RNA-seq imputation based on deep transfer learning using bulk cell information

**DOI:** 10.1101/2024.03.20.585898

**Authors:** Liuyang Zhao, Jun Tian, Yufeng Xie, Landu Jiang, Jianhao Huang, Haoran Xie, Dian Zhang

**Author notes:** The first two authors contributed equally to this paper.

## Abstract

**Motivation:** The growing amount of single-cell RNA sequencing (scRNA-seq) data allows researchers to investigate cellular heterogeneity and gene expression profiles, providing a high-resolution view of transcriptome at the single-cell level. However, dropout events, which are often present in scRNA-seq data, remain challenges for downstream analysis. Although a number of studies have been developed to recover single-cell expression profiles, their performance is sometimes limited by not fully utilizing the inherent relations between genes.

**Results:** To address the issue, we propose a deep transfer learning based approach called scDTL for scRNA-seq data imputation by exploring the bulk RNA-sequencing information. scDTL firstly trains an imputation model for bulk RNA-seq data using a denoising autoencoder (DAE). We then apply a domain adaptation architecture that builds a mapping between bulk gene and single-cell gene domains, which transfers the knowledge learned by the bulk imputation model to scRNA-seq learning task. In addition, scDTL employs a parallel operation with a 1D U-Net denoising model to provide gene representations of varying granularity, capturing both coarse and fine features of the scRNA-seq data. At the final step, we use the cross-channel attention mechanism to fuse the features learned from the transferred bulk imputer and U-Net model. In the evaluation, we conduct extensive experiments to demonstrate that scDTL based approach could outperform other state-of-the-art methods in the quantitative comparison and downstream analyses.

## 1 Introduction

Single-cell RNA sequencing (scRNA-seq) technology has significantly advanced the study of cellular heterogeneity and enabled the high-resolution gene expression profiling at the single-cell level. However, scRNA-seq data always suffers from a high ratio of “dropouts” due to technical limits or improper sequencing. This issue leads to genes not being adequately observed, resulting in artificial zeros in the expression matrix. Such dropout events have been the major obstacle in further downstream analysis such as cell-type clustering [1], trajectory analysis, and differential expressed gene analysis. Therefore, it is desirable to develop an effective method that addresses this critical issue to facilitate scRNA-seq data understanding.

Recently, a number of imputation methods have been proposed to estimate the missing or dropout values, thereby enhancing the interpretability of scRNAseq data. MAGIC [2], one of the earliest available solutions, recovers incomplete single-cell gene expression. It uses the shared information to estimate the missing values in the expression matrix by considering similar cells based on data diffusion. Besides, scImpute [3] conducts separate regression model for each cell to directly estimate true gene expression levels. To measure the uncertainty of estimated values, SAVER [3] uses a Bayesian model with a Poisson LASSO regression and takes advantage of other genes as predictors. DrImpute [4] and KNN smoothing [5] impute scRNA-seq data by averaging or smoothing the expression values via cell clustering to identify similar cells (e.g., KNN). Since the clustering conditions in different datasets are usually unknown, the results heavily rely on similar-cell information that can not be guaranteed.

Instead of using computational methods only, deep learning-based imputation approaches have been effectively applied and achieved satisfactory results. Studies such as DeepImpute and scScope [6, 7] use encoder as a powerful feature extractor that aggregates the cell representation, and reconstructs the expression profile of scRNA-seq data from the latent space with a decoder. Moreover, researchers have extended basic FNN architecture to graph-based models to learn representations of the graph topology and build cell-to-cell similarity links. For example, GraphSCI [8] estimates missing expression values from scRNA-seq data by combining graph convolution and autoencoder neural networks. scGNN [9] introduces a multi-modal framework that adopts dense and graph convolution networks revealing heterogeneous gene expression patterns. Deep neural networks often can better handle scRNA-seq data, but the expression of a gene is often overregulated by others and the model may learn unreliable cell features due to the discrepancy of cell relationship based on dropout noise data [10].

More recently, contrastive learning is also emerged as a promising approach for scRNA-seq analysis [11, 12]. scGCL [13] leverages graph contrastive learning to capture the topological information of the cell graph. It trains a ZINB autoencoder to reconstruct the single-cell expression by randomly dropping out nodes and edges of the cell relationship graph. In addition, CL-Impute [11] uses contrastive learning and self-attention network that captures latent biological cell relationships. To learn accurate cell representations, it generates augmented cells for each original cell and randomly masks non-zero values to simulate dropout events. However, most of aforementioned methods only consider single-cell RNA-seq (e.g., similar cells) information based on the reconstruction loss or contrastive loss (similarity). The inherent relationships between genes are still not comprehensively explored [14], resulting in limited imputation performance - a number of dropout events are still not adequately recovered.

On the other hand, bulk RNA sequencing measures the average expression across hundreds to millions of cells. It could help us to get a global idea of gene expression and relationship between different samples. [14] builds a gene relation network using single-cell and bulk genomics information to facilitate scRNA-seq data imputation. It could yield more accurate results by considering the inherent relations between genes. What is more, [15] proposed SCRABBLE, a matrix regularization based framework that uses bulk cell data as a constraint recovering dropout values in scRNA-seq. Though SCRABBLE does not impose any specific statistical distribution for gene expression levels, it reuiqres the collection of matched bulk data on the same cell/tissue, and consistent cell population is also needed due to matrix regularization for scRNA-seq imputing. [16] developed scDEAL that leverages transferred model on bulk cell data to facilitate single-cell drug response prediction. Therefore, these methods well examined the capabilities of transferring valuable bulk-level knowledge to the single-cell level tasks.

Inspired by above approaches, we introduce scDTL, a deep transfer learning based framework to recover dropout events in scRNA-seq data by fully exploring the bulk cell information. scDTL is capable of dealing with single-cell and bulk from different tissues, and does not force a consistent cell population. Specifically, we first utilize an autoencoder network that learns bulk cell representations with compression and reconstruction scheme. At next step, scDTL harmonizes bulk and single-cell embeddings to ensure that the imputation model is transferable for scRNA-seq learning tasks. Besides, we further explore the scRNA-seq gene representations with different granularity by utilizing a 1D U-Net denoising model as a parallel operation. The U-net model allows the network to propagate contextual information to higher-resolution layers that help capture both coarse and fine features of the scRNA-seq data. Finally, in order to fuse the scRNA-seq features learned from the transferred bulk imputer and U-Net model, we employ a cross-channel attention mechanism on scRNA-seq embeddings. The proposed attention module not only exploits channel-wise attention that better address the heterogeneity of correlations between two output streams, but also focuses on selecting more meaningful spatial features for scRNA-seq representations. Both reconstruction loss and cell clustering loss of gene expression are included in each training epoch.

In the evaluation, we conduct extensive experiments based on six scRNA-seq datasets and use six well-known state-of-the-art imputation methods as baselines. The results demonstrate that scDTL based approach could outperform other solutions in the quantitative comparison including Pearson correlation coefficients (PCCs), Root mean square error (RMSE) and L1-distance, as well as in downstream analyses including cell clustering [17] and pseudotime analysis [18, 19]. We also provide an ablation study to examine the effectiveness of hyper-parameter and modules in scDTL model.

## 2 Materials and Methods

We introduce scDTL, a deep transfer learning based framework that addresses single-cell RNA-seq imputation problem by considering large-scale bulk RNA-seq information synchronously. scDTL mainly consists of two parts: 1. build a imputation model via supervised learning using large-scale bulk RNA-seq data, and 2. propose a framework leveraging well-trained bulk imputation model and a 1D U-net module for imputing the dropouts of a given single-cell RNA-seq expression matrix. Additionally, we also applied an cross-channel attention mechanism which can combine both bulk RNA-seq information and scRNA-seq information and capture their relationships from the global perspective.

The whole architecture of scDTL is illustrated in Fig. 1, there are four components including data preparation and preprocessing, bulk RNA-seq imputation, DTL training and single-cell RNA-seq imputation.

**Fig. 1.**
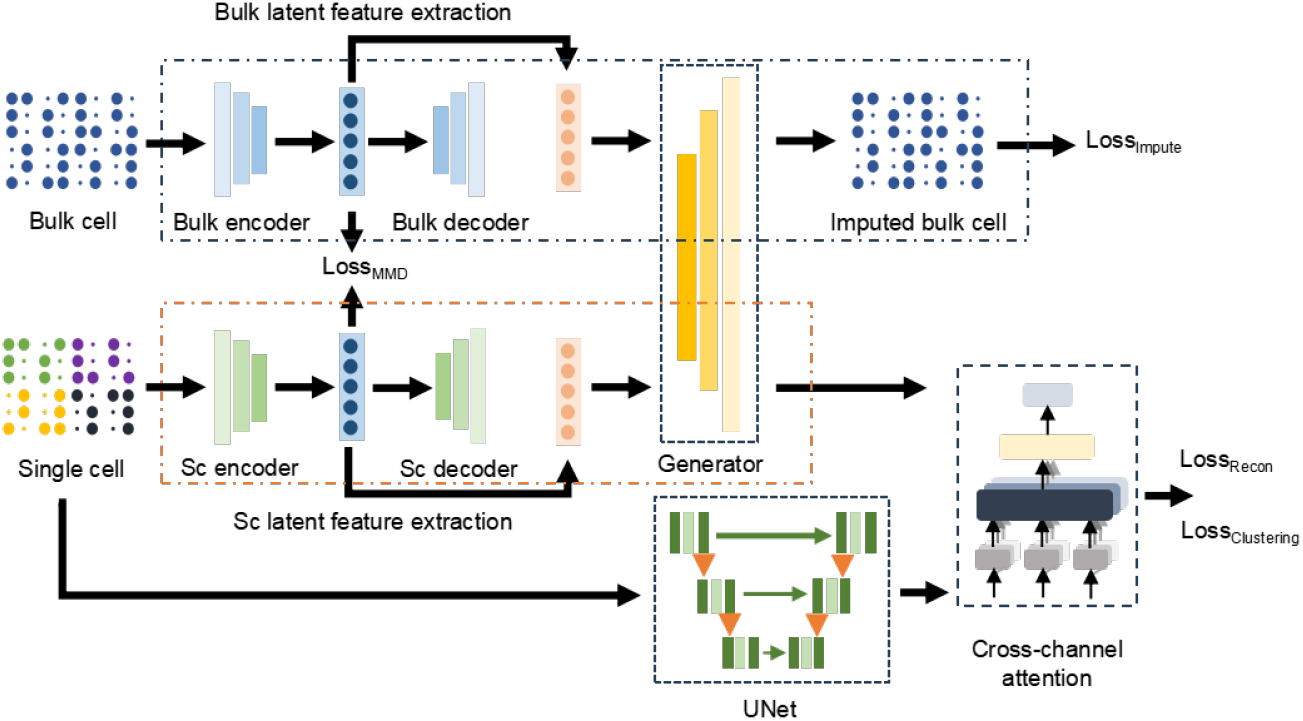
The workflow of scDTL. (i) Bulk RNA-seq imputation training, (ii) RNA-seq imputation DTL training, and (iii) Single-cell RNA-seq imputation training.

### 2.1 Data Preparation and Preprocessing

In this study, six single-cell RNA-seq datasets of tumor cells [20–22] and one CCLE bulk RNA-seq dataset [23] were adopted to examine the ability of scDTL in dropout events imputation. The detailed information of datasets are summarized in Table 1. As we can see from the table, the number of genes ranges from 12274 to 18120, the number of cells ranges from 507 to 24427,the number of cell clusters (groundtruth label) varies from 2 to 10, and the median of zero-value proportion 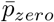 is in 0.431−0.937.

**Table 1.** Summary of the scRNA-seq datasets.

| Dataset | Cell | Gene | Cell type | $\bar{p}_{zero}$ | Platform |
| --- | --- | --- | --- | --- | --- |
| GSE112274 | 507 | 13801 | 5 | 0.546 | Drop-seq |
| GSE117872 | 1302 | 18120 | 10 | 0.555 | 10x |
| GSE134836 | 24427 | 14467 | 2 | 0.867 | CEL-seq2 |
| GSE134838 | 8112 | 12274 | 2 | 0.937 | inDrop |
| GSE134839 | 2903 | 12811 | 6 | 0.891 | Smart-seq2 |
| GSE134841 | 5662 | 12942 | 5 | 0.915 | Drop-seq |

In order to reduce the technical variance and control the sample quality in each scRNA-seq dataset, data preprocessing were performed. Due to the high dropout rate of scRNA-seq expression data [16, 9], we firstly filtered out genes expressed as non-zero in less than 1% of cells, and cells expressed as non-zero in less than 1% of genes. In addition, the gene matrices 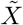 of bulk and single-cell were normalized by dividing by the total UMI count (refer to Python package SCANPY [24]), multiplied by factor = 10^6^.

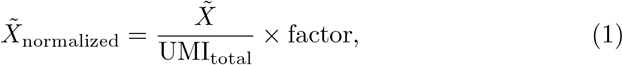

Then we performs a log transformation (log1p) on the normalized counts. The addition of 1 helps avoid issues with taking the logarithm of zero.

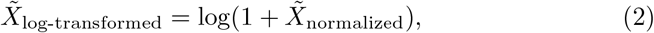

The values in *X*∼_log-transformed_ are then scaled (from 0 to 1) using Min-Max scaling:

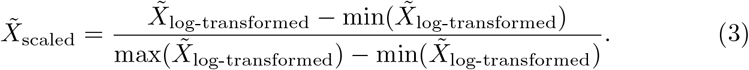

we select the top 4000 highly variable genes into gene expression matrix *X*_*b*_ and *X*_*sc*_ [25]. We split both bulk RNA-seq data and single-cell RNA-seq data 64%, 16%, and 20% as the training set, validation set, and testing set, respectively.

### 2.2 Bulk RNA-seq Imputation Training

Firstly, we build a bulk RNA-seq imputation model using a denoising autoencoder (DAE). The DAE is applied to learn the low-dimensional representation of the bulk expression matrix *X*_*b*_, which is derived from section 2.1. The DAE takes the input *X*_*b*_ and intentionally corrupts it by adding noise - randomly masking (replace non-zero RNA expression values with zero). Then an encoder-decoder model is trained to reconstruct the original bulk RNA-seq expression from the corrupted one. The training workflow is composed of three parts:

1. **Encoder:** The goal is for the autoencoder to learn a representation that is robust enough to capture the underlying structure of the data and remove the introduced noise during the reconstruction process. In our design, the encoder *E*_*b*_ will learn to capture relevant gene features of bulk RNA-sq, and map the input data with random masking (simulated dropout operations) into a hidden representation as a compressed form.
2. **Decoder/Imputer:** The latent representation is then fed into a decoder neural network *D*_*b*_. The decoder’s task is to reconstruct input from the compressed representation and impute the bulk RNA-seq dropout values.
3. **Reconstructed Output:** The final output *X*_*bi*_ of the DAE is the attempt to reconstruct the original, uncorrupted input *X*_*b*_ without the noise (dropout values). The DAE is optimized by the reconstruction loss function (Mean Squared Error, MSE) - minimizing the difference between the input *X*_*b*_ and the reconstructed output *X*_*bi*_:

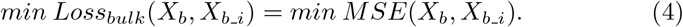

### 2.3 Single-cell RNA-seq Feature Extraction

In order to pretrain an encoder for the extraction of low-dimensional features from the single-cell RNA-seq data, we use a similar DAE model as in the previous section 2.2. We add noise *n*_*s*_ to the single-cell RNA expression *X*_*sc*_ based on a binomial distribution for generating 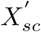.

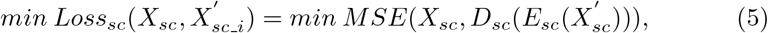

where *E*_*sc*_ and *D*_*sc*_ are the encoder and decoder for the noisy single-cell RNA expression input 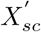, and the reconstructed result 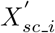 is the output of the decoder *D*_*sc*_. In the testing phase, we will also use the input data with simulated dropout events for *E*_*sc*_ to examine the imputation performance.

### 2.4 RNA-seq Imputation DTL Training

By leveraging the knowledge derived from the Section 2.2, we are able to enhance single-cell RNA-seq imputation by training the model that learns general features and representations from the source bulk RNA-seq data. More specifically, we apply a domain adaptation architecture inspired by Domain Adversarial Neural Network (DANN) [26]. The proposed architecture promotes the emergence of features that are both discriminative and invariant to the change of domains (from bulk RNA-seq to single-cell RNA-seq).

As the training progresses, we update the both two encoders *E*_*b*_ and *E*_*sc*_ to balance the distribution of features extracted from bulk and single-cell domains by introducing the MMD (Maximum Mean Discrepancy) loss. The MMD loss is used to measure the similarity between the outputs of *E*_*b*_ and *E*_*sc*_, which is often defined as the squared difference between the empirical mean embeddings of the source and target domains in a reproducing kernel Hilbert space (RKHS) and can be expressed as:

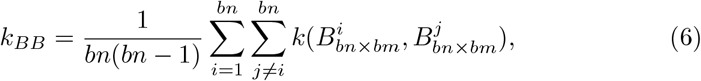

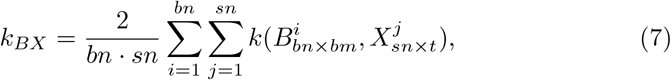

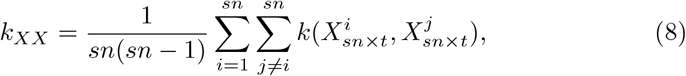

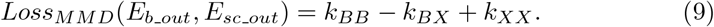

where *k*(*x*_*i*_, *x*_*j*_) is the kernel function applied to samples from the source domain, and *k*(*y*_*i*_, *y*_*j*_) is the kernel function applied to samples from the target domain. In addition, we also condiser the bulk imputation loss (derived from the imputer *D*_*b*_) together with the the MMD loss during the whole training process as defined below:

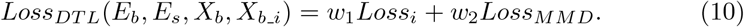

Where *w*_1_ and *w*_2_ are the weights of *Loss*_*i*_ and *Loss*_*MMD*_.

By joinly optimizing *E*_*b*_, *E*_*sc*_, and *D*_*b*_, the proposed domain adaptation architecture is able to minimize the distribution discrepancy between the source and target domains in the RKHS, effectively aligning bulk and single-cell feature distributions.

### 2.5 Single-cell RNA-seq Imputation Training

Using the well-trained encoder *E*_*sc*_, we can now compress any single-cell RNA-seq data into a compact and low-dimensional latent variable *z*_*sc*_ = *E*_*sc*_(*X*_*sc*_). Then the transferred bulk imputer *D*_*sc*_ will take the latent variable *z*_*sc*_ as input, while at the same time a 1D U-Net denoising model is applied to extract and compute single-cell RNA-seq features at multiple scales simultaneously. Given two outputs *X*_*scDTLi*_ and *X*_*scUi*_, we leverage a cross-channel attention module to sequentially infer attention maps along two separate channels and dimensions (both bulk imputer and single-cell U-Net).

#### 1D U-net architecture

Unlike the FCN-based decoder that complements the typical contracting network with continuous layers, the U-Net features M levels of downsampling and upsampling blocks replacing for increasing the resolution of the output. In our design, each block with contracting path follows a typical structure of 1D convolutional network. It consists of the repeat application of one 1 × 1 conv layer with stride 2 (unpadded), one 3 × 1 conv layer followed by a Sigmoid function with stride 2 (padding = 1). Moreover, we double channel numbers at each downsampling step (16-32-64-128), while at each upsampling step halve the channel numbers with a 3 × 1 conv kernel and concat with the correspondingly cropped feature map from the contracting path. At the final step, one 1×1 conv layer is used to map each 16 component feature vector to the desired number of gene expression values (4000 in our case). There are in total 23 conv layers in the proposed U-net structure.

#### Cross-channel attention module

As shown in Figure 2, given a concatenation *X*_*att*_∈ *R*^*C*×*L*^ of *X*_*scDTLi*_ and *X*_*scUi*_ as input, the cross-channel attention calculates the channel and spatial weight matrix sequentially, selecting ‘what’ and ‘where’ is meaningful in gene expression analysis. The overall process is summarized as follows:

**Fig. 2.**
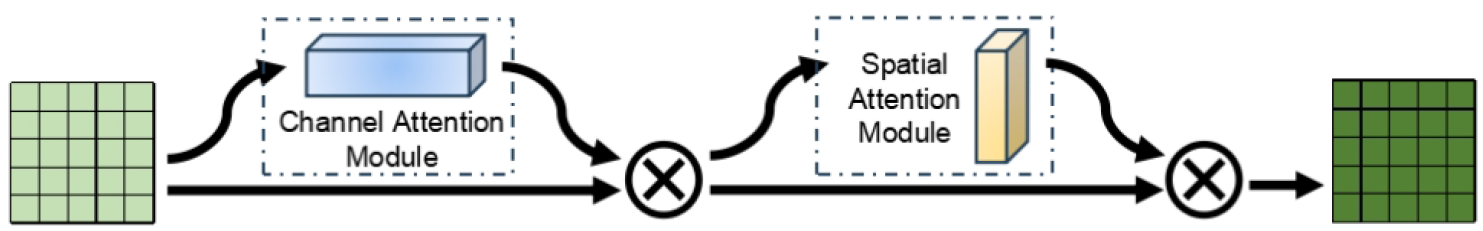
The architecture of cross-channel attention module.

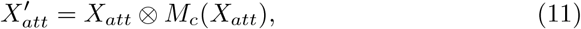

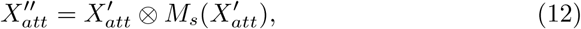

where *M*_*c*_ ∈ *R*^*C*×1^ is the 1D channel attention matrix, *M*_*s*_ ∈ *R*^1×*L*^ is the 1D spatial attention matrix, and ⊗ denotes element-wise multiplication. 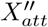 is the final refined output.

To produce a channel attention matrix *M*_*c*_ of feature map *X*_*att*_, we squeeze the spatial dimension of the feature map by applying both average-pooling *P*_*avg*_ and max-pooling *P*_*max*_. It has been empirically examined that exploiting both average-pooled and max-pooled features could greatly improve the representation power of networks rather than using each independently [27].

The pooling features are connected by a shared convolutional network *ξ*, which is usually composed of multi-layer perceptron (MLP) with one hidden layer:

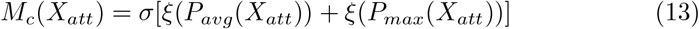

where σ represents the Sigmoid function.

To emphasize the inter spatial features of gene expression, the spatial attention refines the feature map along channel dimension by leveraging average pooling and max pooling operations to concatenate an efficient feature descriptor. The spatial attention map can be generated by applying a two channel convolution layer as shown below:

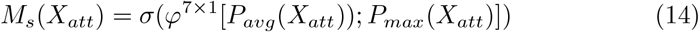

where *φ* is the convolutial function with a (7, 1) kernel size. For detailed explanation and discussion on channel-wise attention mechanism, we refere to the paper [27].

Finally, we integrates the cell-clustering loss with weight *β* and gene reconstruction loss together to retain the single-cell heterogeneity during the training process. The well-trained *E*_*sc*_ and imputer *D*_*b*_ together with the U-net module will take the single-cell RNA-seq data *X*_*sci*_ as the input, and output the imputed gene expression *X*_*atti*_ by replacing the zero values in cell *X*_*sci*_.

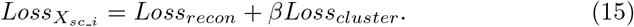

## 3 Results

### 3.1 Experimental Settings and Baselines

In the experiment, we regarded the gene expression from six single-cell datasets as ground-truth. We simulate the dropout events in selected single-cell testing sets by randomly masking 10%, 20% and 40% of non-zero gene values (replacing the value with zero). We then performed the imputation task on both synthetic dropout data and original data with zero values using scDTL and the baseline methods to recover the missing values. Next, we quantitatively evaluated the performance of each method on the synthetic dropout data by using benchmarking metrics that measure the similarity and error distance between the recovered data and the ground-truth data. We also provided downstream analysis tasks to evaluate the performance of each method on the original data with zero values. Downstream tasks include cell clustering [17], PAGA (Partition-based Graph Abstraction) [18] and pseudotime analysis [19].

For comparison, we selected six state-of-the-art methods for single-cell RNA-seq data imputation as baselines including MAGIC [2], CMF-Impute [28], GE-Impute [29], CL-Impute [11], scGCL [13] and SCRABBLE [15]. All parameters and configurations of baseline methods in the comparison follow the settings as suggested in the original papers or the default parameters of shared codes.

### 3.2 scDTL greatly improved the performance of missing data recovery

To assess the performance of scDTL in imputing missing values in scRNA-seq data, we simulated dropout events by randomly masking 10%, 20%, and 40% of non-zero expression values in six datasets. Then we adopted three quantitative metrics including Pearson correlation coefficients (PCCs), Root mean square error (RMSE) and L1-distance to evaluate the accuracy of the imputed values. The formulations of three quantitative metrics are shown as follows:

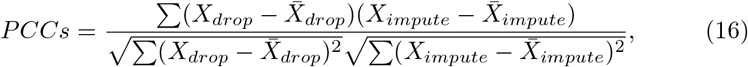

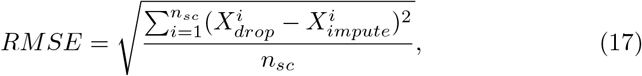

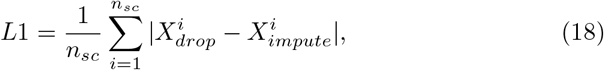

where 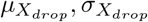 represent the mean and variance of *X*_*drop*_ and 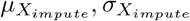 represent the mean and variance of *X*_*impute*_, and *n*_*sc*_ is the total number of the single-cell samples.

As shown in Fig. 3 and Supplementary Table 1, scDTL outperformed other methods in nearly all cases of 10% and 20% dropout rates with higher PCCs values as well as lower RMSE and L1-distance values. Notably, scDTL showed excellent performance in recovering the missing values at 40% dropout rates than any other methods in all datasets. These data indicate that our model excels at precisely recovering missing data, especially in situations with high dropout rates. We argue that the scDTL deep transfer learning framework takes advantage of the original gene expression values from bulk data, rather than simply calculating cell-to-cell similarity in scRNA-seq data. This is crucial, especially when ‘real’ similar neighbors may not be collectible due to a high ratio of dropouts in scRNA-seq data. Although SCRABBLE also leverages bulk cell information, scDTL employs neural network-based methods with strong feature extraction capabilities. This allows the model to learn latent representations effectively, even in the presence of incomplete gene expressions. Overall, scDTL effectively recovers missing values in scRNAseq data, producing an imputed matrix that closely resembles the actual data.

**Fig. 3.**
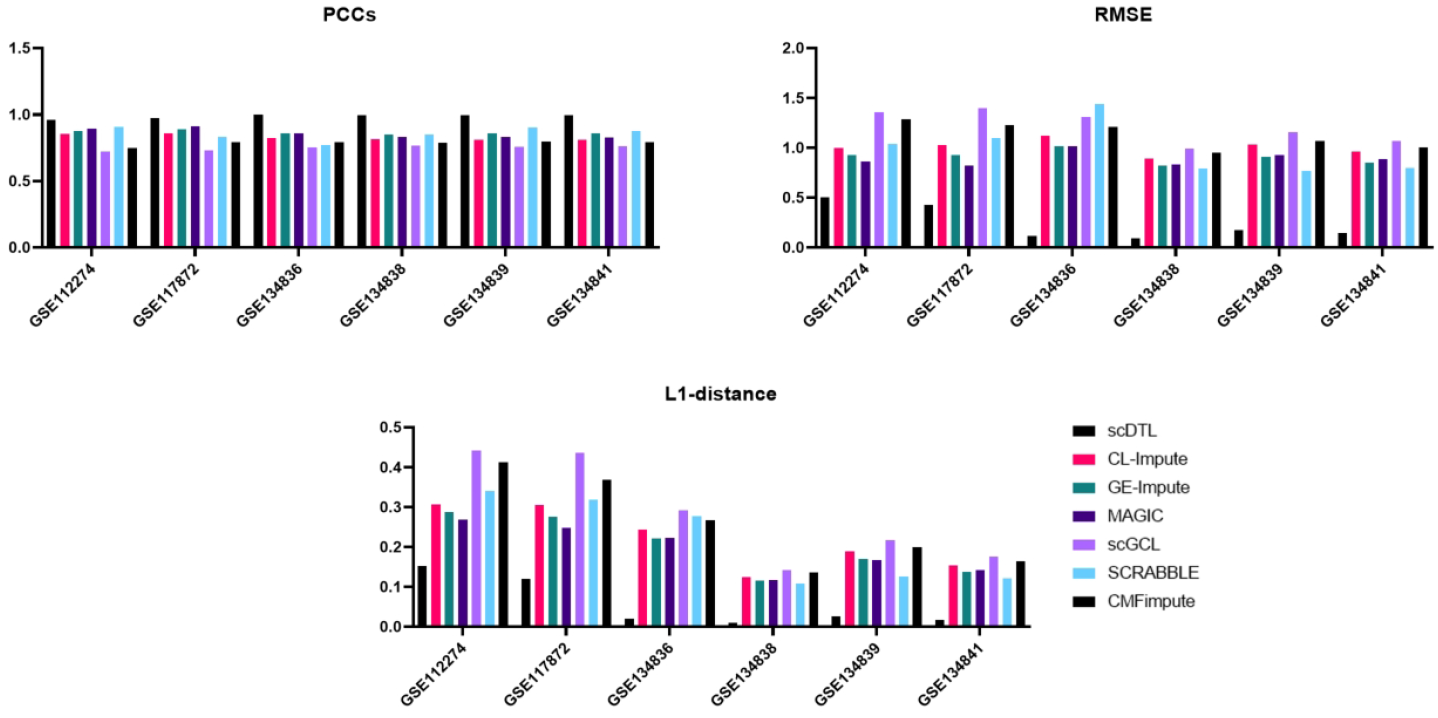
Quantitative metrics of scDTL compared with other imputation methods in recovering missing values in scRNA-seq data at 40% dropout rates

### 3.3 scDTL significantly improved the performance of cell clustering

Cell clustering is a crucial task in scRNA-seq downstream analysis, impacting cell type annotation. To assess scDTL’s clustering performance, we compared it with six baseline methods across six datasets. The clustering was conducted through the SCANPY toolkit [24], and we computed the Adjusted Rand Index (ARI) and Fowlkes-Mallows Index (FMI) to measure the correlation between the clustering results and the annotated cell populations in the datasets [30].

As shown in Fig. 4a, scDTL demonstrated greater improvement in clustering accuracy compared to all other baseline methods in the GSE117872 and GSE134839 datasets, as indicated by higher ARI and FMI values. In datasets GSE134836, GSE134838, and GSE134841, scDTL’s performance was comparable to that of CL-Impute, MAGIC, scGCL, or CMFImpute, and it outperformed the other baseline methods. In dataset GSE112274, scDTL outperformed other methods except for SCRABBLE. We argue that this is because both scDTL and SCRABBLE utilize bulk RNA-seq information for the imputation. However, the limited cell number (500) in this dataset may have restricted scDTL’s capacity to learn latent representations effectively, thereby impacting its performance in downstream analysis. Together, these data suggest that scDTL performed either better or similarly in terms of improving clustering accuracy compared to other methods, and therefore can be applied in a broader range of scenarios.

**Fig. 4.**
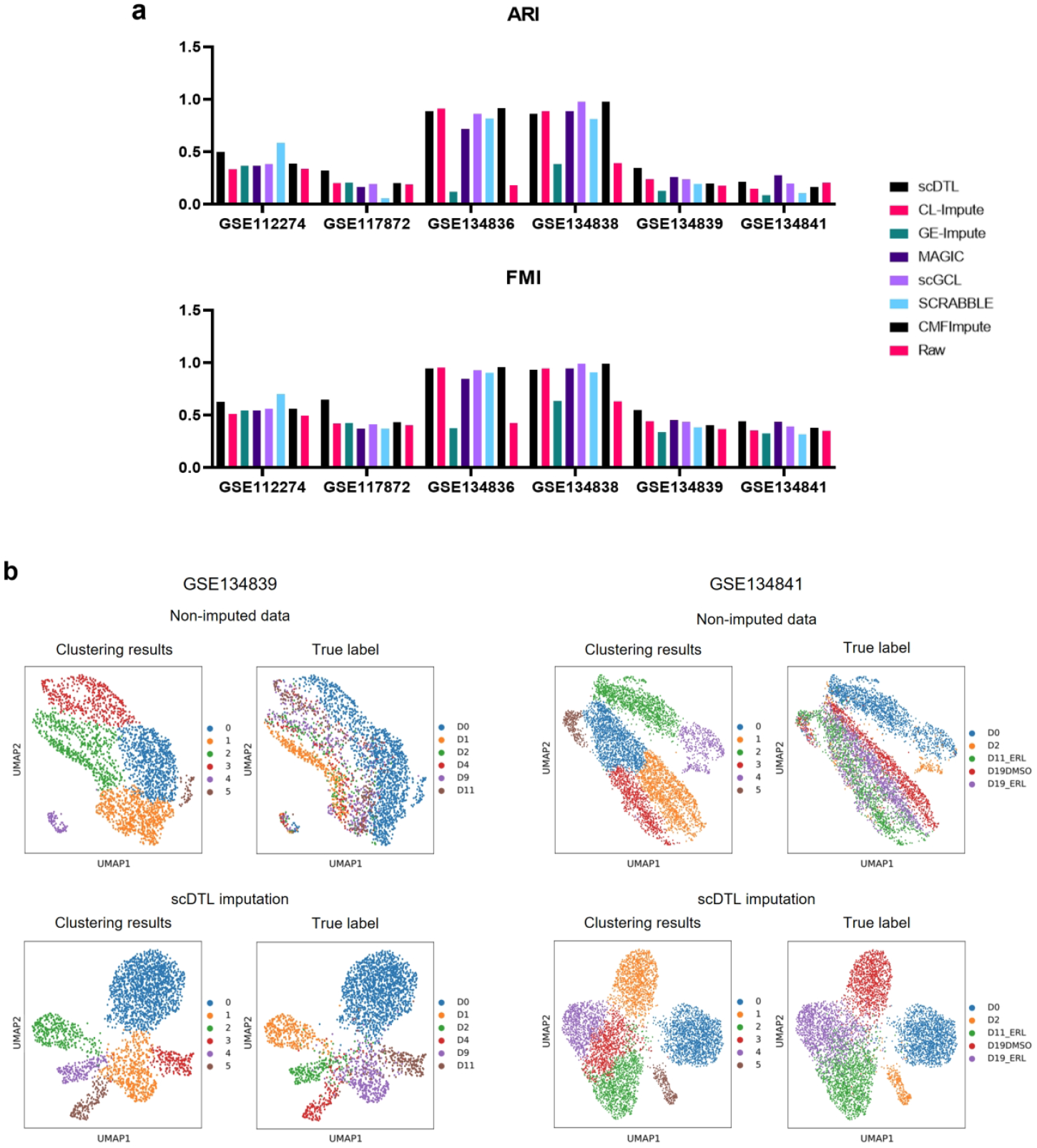
Clustering analysis. a, ARI and FMI values of scDTL clustering results compared with other imputation methods. b, UMAP plots of clustering results in GSE134839 and GSE134841 datasets before and after scDTL imputation.

The clustering results were visualized using the uniform manifold approximation and projection (UMAP) method. As shown in Fig. 4b, for the non-imputed data in the GSE134839 and GSE134841 datasets, the clustering results were completely inconsistent with the true labels. In contrast, in the scDTL imputation plot for the GSE134839 dataset, different clusters were well-separated and consistent with the true labels. Similarly, in the GSE134841 dataset, although unsupervised clustering analysis revealed six clusters compared to the actual five true label clusters, the alignment between the clustering results and true labels was still greatly improved compared to the analysis with non-imputed data. In comparison, the unsupervised clustering results were still inconsistent with the true labels after the imputation using other methods (Supplementary figure 1). These data demonstrate that the clustering results of scDTL are highly consistent with the ground-truth labels.

### 3.4 scDTL improved the performance of trajectory analysis

Trajectory analysis of scRNA-seq data is an important downstream analysis revealing cellular developmental patterns. Dropout events can impact trajectory reconstruction, but imputation methods effectively mitigate this issue. To assess scDTL’s effectiveness in improving trajectory analysis, we reconstructed the cell trajectory imputed by each method. We employed scanpy.tl.paga and scanpy.tl.dpt toolkits to reconstruct trajectory inference for the GSE112274 and GSE134838 datasets, featuring cancer cells treated with tyrosine kinase inhibitor or vemurafenib, respectively, in a time-course manner. We first calculated Pseudo-temporal Ordering Score (POS) [31] to quantify the accuracy of cell pseudo-time ordering compared to the ground-truth cell ordering:

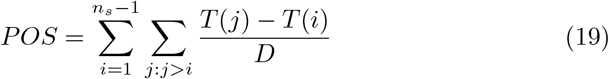

where *T* (*i*), *T* (*j*) is the true-time of *i*th and *j*th cells in cell pseudo-time ordering respectively.

Quantitative assessment using the POS index shows that scDTL achieves the highest consistency between cell pseudo-time and true time in both datasets compared to other methods (Fig. 5a), indicating improved accuracy in inferred cell pseudo-time ordering. Furthermore, as shown in Fig. 5b, scDTL imputed data trajectories align better with the true-labeled time points in comparison with the non-imputed data. These findings highlight that scDTL enhances the precision of inferred cell pseudo-time ordering.

**Fig. 5.**
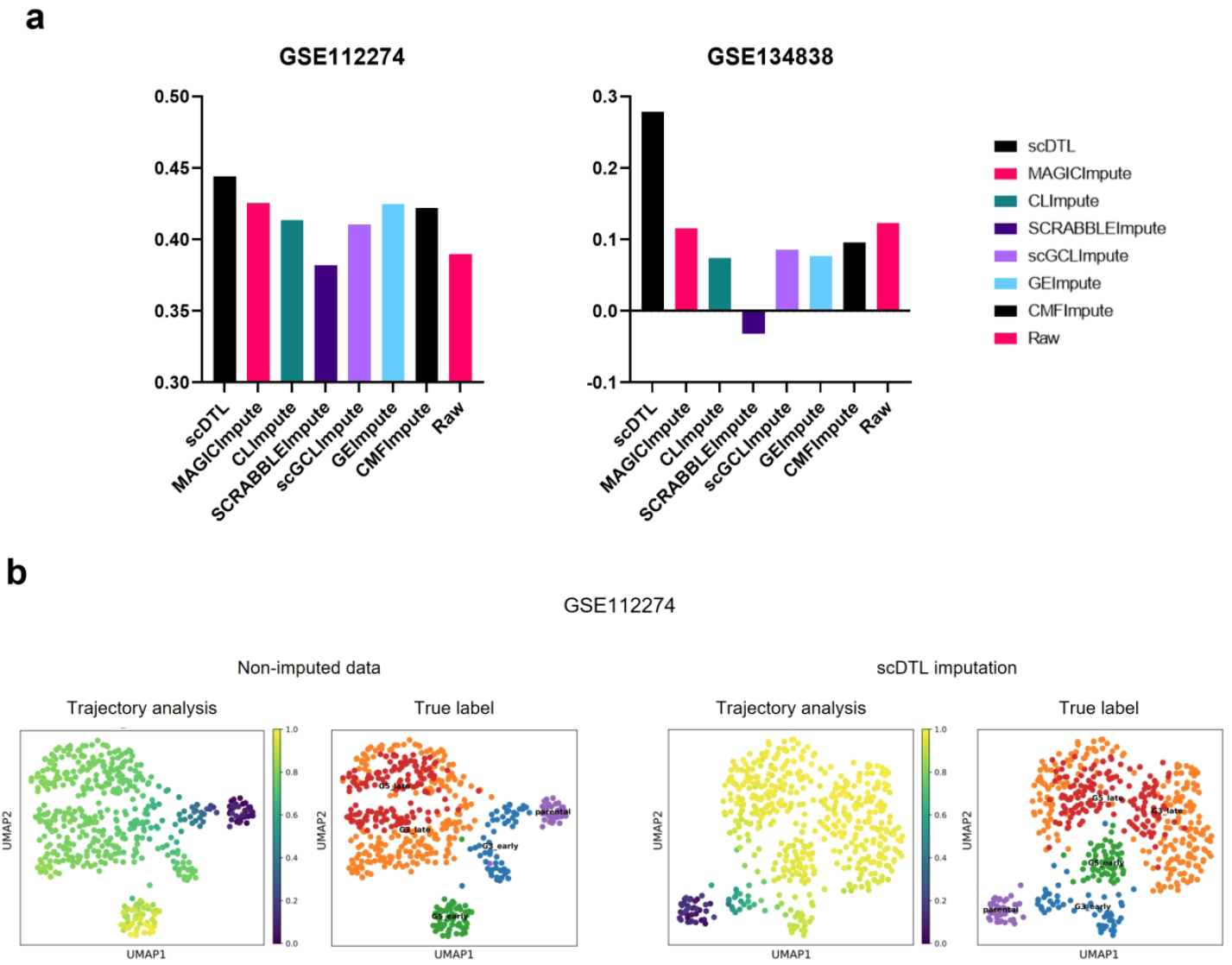
Trajectory analysis. a, POS of the indicated datasets imputed by different methods. b, Visualization of trajectory inference. In true label, G3 and G5 refer to two independent TKI-resistant cell clones. Early and late refer to cells collected at 30 days and 120 days after the treatment.

### 3.5 Ablation Study

To assess the impact of each module on the imputation results, we ablated Unet module and CBAM module when calculating quantitative metrics and clustering metrics in all datasets after the imputation. As shown in Supplementary Table 2, removing either of these modules dramatically decreased PCCs values and increased both RMSE and L1-distance values at the 40% dropout rate. In terms of clustering accuracy, ARI and FMI values also markedly dropped after these components were eliminated, as shown in Supplementary Table 3. These data suggest that Unet module and CBAM module are essential components of the scDTL method.

## 4. Discussion

Single-cell RNA sequencing has significantly facilitated the comprehensive studies of gene expression profiles at the single cell level. Especially in the field of cancer research, scRNA-seq has been applied to investigate the cellular heterogeneity in the tumor microenvironment and tumor cells themselves. However, dropout events have become a major challenge that significantly diminished the accuracy of scRNA sequencing analysis. Consequently, it is imperative to develop effective imputation methods to recover the incomplete gene expression profiles. In this article, we used scRNA-seq data and bulk RNA-seq data from tumor cells to examine the performance of a novel gene-imputation framework - scDTL as a proof-of concept. The evaluation results shows that the proposed method can be further applied in different cell types or tissues, thereby enabling a broader range of research scenarios such as cancer research and other diseases.

Typically, there is always a high ratio of ‘dropouts’ in scRNA-seq matrices (can contain up to 90% zero values), while there are 15%−40% of genes in bulk RNAseq of various tissues are not expressed. Unlike existing imputation methods that only condiser scRNA-seq data taking advantage of the similarity among cells and genes. We aim to explore inherent relationships between genes by leveraging bulk cell data. Specifically, scDTL applies a domain adaptation technique that preliminarily trains the imputation model for scRNA-seq data using a large volume of bulk RNA-seq data. We also employ a parallel operation with a cross-attention mechanism to balance the feasibility of harmonizing scRNA-seq data with bulk data while preserving original single-cell data information.

In the evaluation, we examine the imputation performance of scDTL and other state-of-the-art baseline methods by conducting extensive experiments including quantitative metrics, clustering analysis, PAGA (Partition-based Graph Abstraction) and pseudotime analysis. The results demonstrate that scDTL could outperform other solutions in most cases on scRNA-seq 6 datasets. In the future, we would like to continue to enhance the capability of scDTL and use unsupervised learning method such as contrastive learning. As it allows the model to gain a self-perspective in exploring cell representations, we can capture and learn the scRNA features without labeled data.

## 5 Availability and Implementation

The code and data of scDTL are available at: https://github.com/bleedingseraphY/scDTL.git

