## Supplementary figure 1 for "scDTL: single-cell RNA-seq imputation based on deep transfer learning using bulk cell information"

### Slide 1
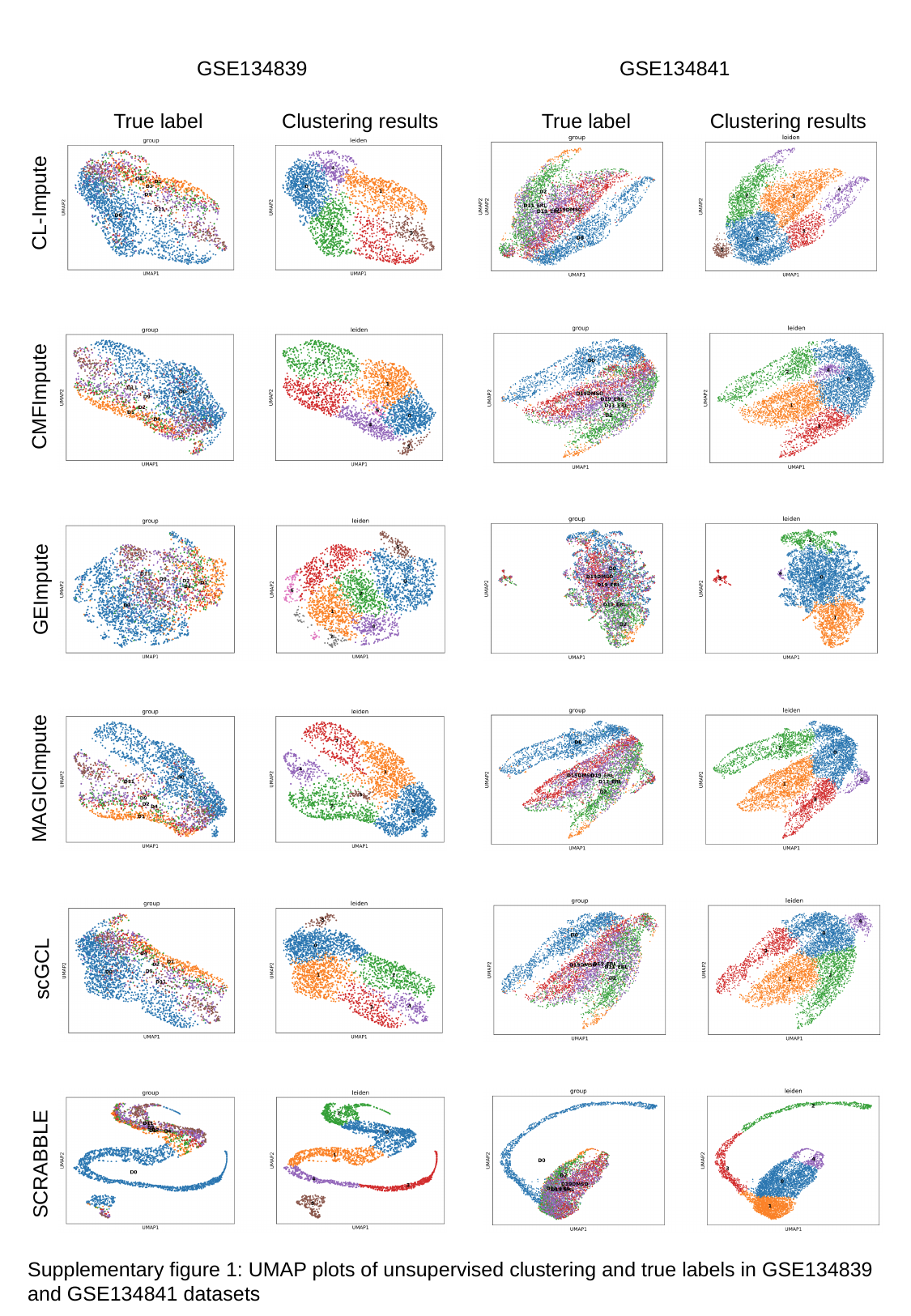

GSE134839
GSE134841
True label Clustering results
True label Clustering results
CL-Impute
CMFImpute
GEImpute
MAGICImpute
scGCL
SCRABBLE
Supplementary figure 1: UMAP plots of unsupervised clustering and true labels in GSE134839 and GSE134841 datasets
